# RBV: Read balance validator, a tool for prioritising copy number variations in germline conditions

**DOI:** 10.1101/340166

**Authors:** Whitney Whitford, Klaus Lehnert, Russell G. Snell, Jessie C. Jacobsen

**Affiliations:** School of Biological Sciences, The University of Auckland, New Zealand; Centre for Brain Research, The University of Auckland, New Zealand

**Author notes:** Corresponding author: Whitney Whitford, University of Auckland, School of Biological Sciences, Private Bag 92019, Auckland 1142, New Zealand.

**Keywords:** Copy number variation, read balance, allele specific copy number, genomics, molecular diagnostics, read balance validator

## Abstract

**Background:** The popularisation and decreased cost of genome resequencing has resulted in an increased use in molecular diagnostics. While there are a number of established and high quality bioinfomatic tools for identifying small genetic variants including single nucleotide variants and indels, currently there is no established standard for the detection of copy number variants (CNVs) from sequence data. The requirement for CNV detection from high throughput sequencing has resulted in the development of a large number of software packages. These tools typically utilise the sequence data characteristics: read depth, split reads, read pairs, and assembly-based techniques. However the additional source of information from read balance, defined as relative proportion of reads of each allele at each position, has been underutilised in the existing applications.

**Results:** We present Read Balance Validator (RBV), a bioinformatic tool which uses read balance for prioritisation and validation of putative CNVs. The software simultaneously interrogates nominated regions for the presence of deletions or multiplications, and can differentiate larger CNVs from diploid regions. Additionally, the utility of RBV to test for inheritance of CNVs is demonstrated in this report.

**Conclusions:** RBV is a CNV validation and prioritisation bioinformatic tool for both genome and exome sequencing available as a python package from https://github.com/whitneywhitford/RBV

## Background

There are four main types of variation in the human genome: single nucleotide variants (SNVs), small scale changes in genomic content in the form of short indels, structural variants, and aneuploidies. Structural variants consist of medium to large scale changes to the genomic structure, and includes both balanced chromosomal rearrangements (such as inversions and translocations) and copy number variants (CNVs). CNVs are typically defined as deletions or multiplications of sections of the genome, resulting in changes of genomic content greater than 1 kb [1]. Initial efforts to map genetic variation on the whole genome scale indicated that SNVs constituted the majority of variation between individuals [2]. However, large scale collaborations mapping CNVs in the human genome found on average an individual harbours over 1,000 CNVs of 443 bp or greater [3–6]. Taken together, although there is a greater number of SNVs per individual (approximately 3.6 million or ~0.1% of the genome [5]), due to the greater average size of CNVs and indels, they are responsible for greater genomic variance between genomes (up to 48.8 Mb or ~1.5% [6]).

CNVs play an important role in gene expression with changes in genetic content larger than 1Mb estimated to be responsible for 17.7% of the genetic impact on gene expression [7]. One would expect that the proportion of genetically controlled variation in gene expression would be higher if CNVs smaller than 1Mb were included in such analyses. CNVs are able to effect gene expression directly through copy number changes of genes and regulatory elements [8], and indirectly through unmasking of recessive alleles [9] and positional effects [10]. As such, there has been increasing volume of research into the role of CNVs in disease. In particular, CNVs have been implicated in the aetiology of neuropsychiatric disorders including schizophrenia, intellectual disability, and autism spectrum disorder (as reviewed by Malhotra & Sebat, 2012 [11]). Therefore, chromosomal microarray (CMA) has become a first-tier clinical diagnostic test for patients with unexplained intellectual disability, autism spectrum disorder, or multiple congenital anomalies, with diagnostic yield of 15-20% (reviewed by Miller, et al., 2010 [12]). The use of high throughput sequencing (HTS) in the form of whole exome sequencing (WES) and whole genome sequencing (WGS) is increasing for diagnostic testing, both due to its decreasing cost and ability to investigate genetic variants without prior hypotheses. HTS based methods offer the potential of identifying SNV, indels and CNVs not detected by current diagnostic CMA thresholds [13] in a single test.

With the rapid implementation of HTS in molecular diagnostics and research, there has been a proliferation in tools for variant identification. There are currently over 80 software packages designed to identify CNVs from WGS alone [14]. These tools predominantly rely on four characteristics of the sequence data: read depth, split reads, read pairs, and assembly-based techniques (reviewed by Zhao, et al., 2013 [15]). As yet underutilised, the allele balance of reads at a position contributes additional data that can also be exploited for CNV variant detection and validation. This ‘read balance’ is computed from relative read coverage of each allele at a given locus. The read balance can provide information regarding the copy number over the region in the form of the allele-specific copy number (ASCN). Positions in diploid regions of the genome are primarily invariant (homozygous) (as demonstrated in Figure 1A). This is represented by a relative read distribution peak about 1. The heterozygous positions (SNVs) are represented by a normal distribution centred on 0.5, with the reads split evenly across the two alleles. A deleted (hemizygous) region should not contain any heterozygous positions, resulting in a distribution peak centred around 1, as depicted in Figure 1B. A triplicated region as represented in Figure 1C, however, is expected to have homozygous SNVs along with the heterozygous SNVs represented by two normal distributions centred on 0.33 and 0.66, indicating that one third of the reads at a given locus include one allele, and two thirds of the reads include the other.

**Figure 1.**
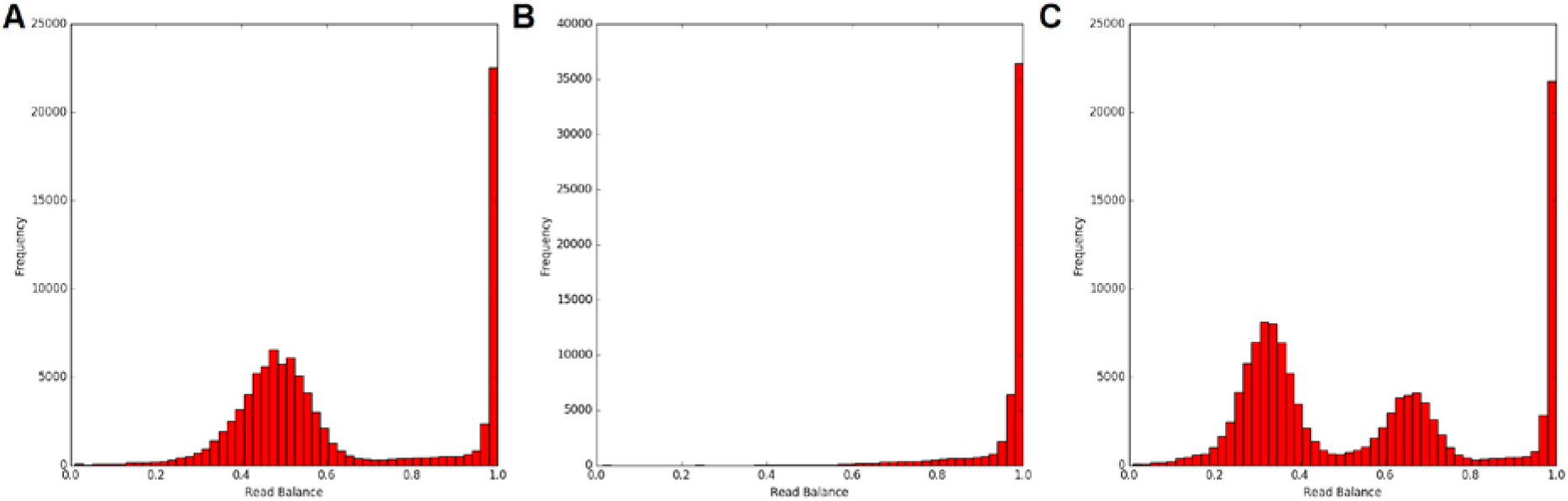
Distribution of relative reads for diploid, haploid, and triploid regions in whole genome sequence. **A.** Expected distribution of all positions in a diploid genome. **B.** Expected distribution of all positions in a hemizygous genome. **C.** Expected distribution of all positions in a triploid genome.

A number of bioinformatic tools have utilised ASCN for determining CNVs in cancer samples [16–24]. These techniques rely on sequence data from paired tumour and normal tissue samples, and therefore are not suitable for identifying germline CNVs. Alternatively, AS-GENSENG[25] and ERDS[26] incorporate read balance information into their algorithms along with read depth based data to discover CNVs. However, there is no independent platform providing validation of CNVs using read balance, allowing for integration of this additional data source in established bioinformatic pipelines that use alternative CNV discovery tools.

RBV utilises read balance data to validate CNVs identified by other software packages, allowing for prioritisation of CNVs for causation in molecular diagnostic testing.

## Implementation

RBV is a python package, which incorporates the read balance data from positions within the CNV of interest with randomly sampled windows across the genome to predict the authenticity of CNVs. The software extracts the read balance information from a variant call format (vcf) file, and uses CNV coordinates from an interval list, and can be employed for both WGS and WES generated data. The analyses can be refined by restricting investigation to callable regions or outside of known gaps in the reference through the inclusion of either an interval list of callable regions, or an interval list of gaps in the reference genome provided by the user. RBV specificity can be adjusted by the user through setting of parameters quality and depth cut-offs at each position in the vcf, readbal cut-off for deletion analyses, and the number of randomly generated permutations for the positions and windows. RBV can incorporate data derived from popular variant callers (HaplotypeCaller[27], SAMtools[28], Freebayes[29], and Platypus[30]), and all aligners. However, issues with read balance calculations may arise from non-uniquely aligned regions of the genome if the aligner of choice places these reads at more than one position in the genome. We therefore recommend using aligners that randomly place reads to only one mappable location by default, such as BWA [31].

RBV is freely available via https://github.com/whitneywhitford/RBV.

## Results

The analysis performed by RBV validates two separate hypotheses: that the putative CNV is a deletion with the region being hemizygous or nullizygous, or that the putative CNV is multiplicated where the region is triploid or greater.

### Deletion analyses

Deletions should represent areas of absence of heterozygosity (AOH), therefore the probability that a deletion exists (p-value) is calculated based on an empirical cumulative distribution function (eCDF). For this calculation, a large number of windows (default 1,000) of the same number of callable base pairs as the CNV of interest are randomly generated and the number of heterozygous SNVs in each window is subsequently calculated. The empirical p-value is calculated using the eCDF (equation 1) for the resulting distribution, with the probability being the proportion of randomly generated windows containing the same number or fewer heterozygous SNVs for the CNV in question.

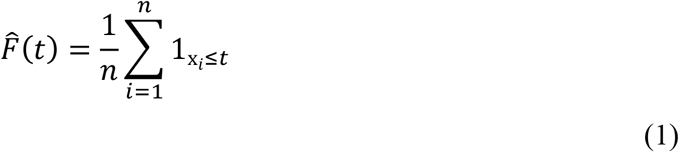

Where *x_1_*, *x_n_* represent *n* number of randomly generated windows of the same size as the CNV, and t is the number of heterozygous SNVs within the CNV of interest.

### Multiplication analyses

The multiplication hypothesis is interrogated using the two-sample Kolmogorov–Smirnov (KS) test, included in the scipy.stats module for Python. For this analysis we only consider the most common allele at each heterozygous position, which gives the distribution in Figure 2A and Figure 2B. The differences in the distribution of read balance for randomly generated diploid heterozygous SNVs and the heterozygous SNVs (default 10,000) in the putative CNV are compared using the 2 sample KS test, represented in Figure 2C.

**Figure 2.**
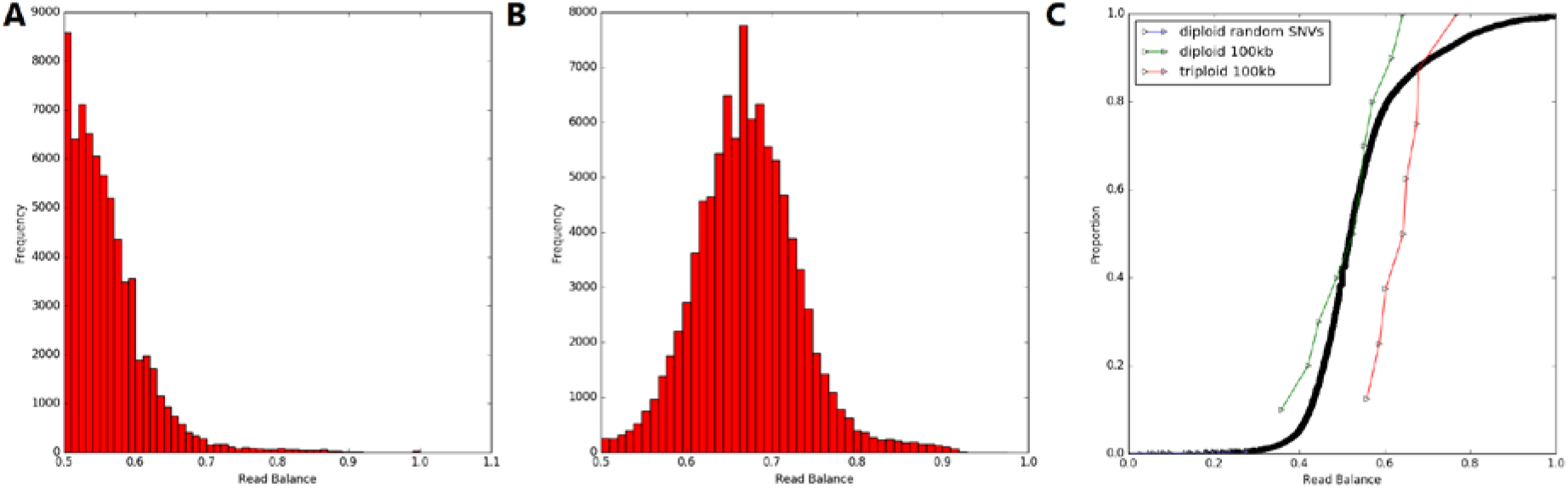
RBV data analysis curves. **A.** Read balance of the most common allele from heterozygous positions in a diploid genome. **B.** Read balance of the most common allele from heterozygous positions in a triploid genome. **C.** CDF curve utilised in a 2-sample KS test, comparing distribution of read balance between randomly generated heterozygous SNVs throughout the reference diploid genome: a 100kb diploid region, and a 100kb triplicated region.

### Performance

To analyse the performance of RBV, a number of common clinically relevant CNVs identified by Matsunami, et al., [32] were simulated using Enhanced Artificial Genome Engine (EAGLE) [33]. Paired regions of the same size and covering the same number of callable positions for each CNV were randomly generated to facilitate comparison. The ability of RBV to prioritise the simulated clinically relevant CNVs over the randomly generated regions is shown in Figure 3. The comparison shows a clear enrichment of the simulated CNVs >10kb with lower p-values, highlighting the performance of RBV with larger CNVs. Therefore RBV will have reduced sensitivity to detect smaller CNVs due to the reliance upon the presence of relatively infrequent heterozygous positions in the randomly generated windows for deletion analysis, and the increased power of a 2-sample KS test with a greater number of heterozygous positions in the CNV. For the regions analysed >10kb, RBV is able to identify statistically significantly (P≥0.05) CNVs with a sensitivity of 0.38 and 1.0 along with a specificity of 1.0 and 0.9, for deletions and duplications (KS), respectively.

**Figure 3.**
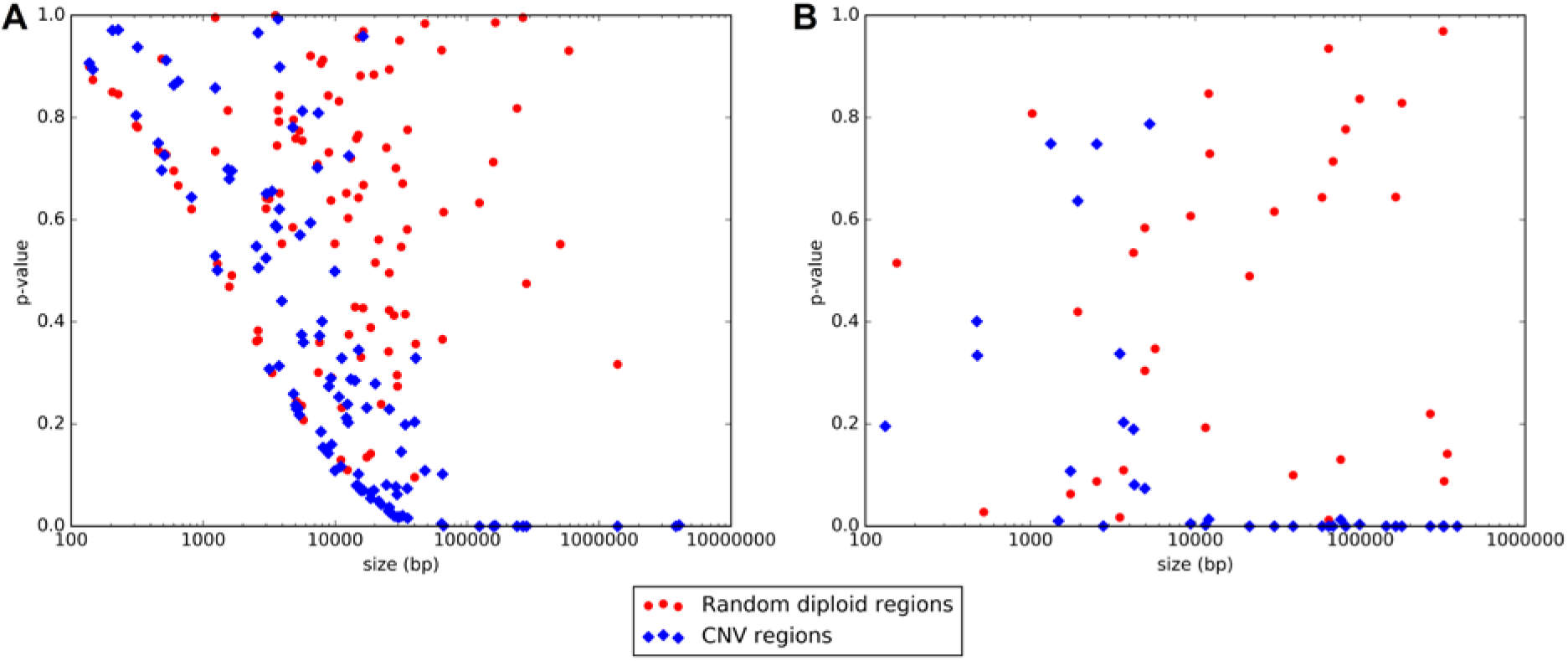
Ability of RBV to prioritise authentic CNVs. Comparision between the results from simulated common causative CNVs identified by Matsunami, et al., [32] and randomly generated diploid regions or the same size and number of callalble positions as each CNV **A.** Performance of RBV for deleted CNVs. **B.** Performance of RBV for duplicated CNVs.

### Use case

Another use for RBV is to test the potential inheritance of CNVs. Using HTS our laboratory recently identified a causative 19.6Mb 2q37 terminal deletion in a child with ASD (reported in [34]). There was both WES and WGS data available for the affected child, and WES data for the parents. RBV was run with default parameters using all four sequence sources, and determined the region (GRCh37 Chr2:233834098-253404903; NC_000002.11:g. 233834098_253404903del) to be deleted with a p-value of 0.0 from both WES and WGS from the affected child (with 0 and 92 heterozygous SNVs predicted in the vcf file over the region, respectively). In comparison, the two parents had 284 and 294 heterozygous SNVs in the exonic sequence in the same region, resulting in p-values of 0.898 and 0.936, respectively. This provided evidence that the causative deletion is *de novo,* which was subsequently confirmed using Sanger sequencing.

## Discussion

As more research and diagnostic centres are investigating the identification of CNVs through sequence data, there is increasing need for the ability to prioritise clinically relevant variants called from the various CNV detection software platforms. Although there are a number of these detection tools that use read depth, split reads, read pairs, and assembly-based techniques, the utility of read balance, or ASCN, in CNV analysis has so far been largely underutilised. Thus, RBV was developed to exploit this additional piece of sequence information to reinforce calls from CNV calling pipelines, allowing for prioritisation of variants in the identification of clinically relevant CNVs.

We compared the results of RBV from simulated clinically relevant CNVs and randomly generated diploid regions. From this we were able to display the ability of RBV to differentiate genuine CNVs >10kb from diploid loci. Thus, this software has utility in highlighting genuine potentially clinically significant CNVs >10kb. However, the utility of RBV decreases for smaller variants. We were also able to demonstrate the ability of RBV to determine the inheritance of a CNV using an example of a clinical case.

## Conclusions

RBV is a software tool designed to assist in the rapidly expanding speciality of identifying clinically relevant CNVs through prioritisation. RBV is a python based package and available under the open source GPL v3 license at https://github.com/whitneywhitford/RBV.

The software includes utility for both multiplication and deletion analysis of nominated CNV sites for both WES and WGS data. Sample data for the operation of RBV is available via the GitHub repository.

## List of abbreviations

AOH: : absence of heterozygosity
ASCN: : Allele-specific copy number
CMA: : Chromosomal microarray
CNV: : Copy number variant
HTS: : High throughput sequencing
kb: : kilo base
RBV: : Read balance validator
SNV: : Single nucleotide variant
WES: : Whole exome sequencing
WGS: : Whole genome sequencing

## Declarations

### Ethics approval and consent to participate

The genetic analysis and de-identified publication of variants was performed under the approval of the New Zealand Northern B Health and Disability Ethics Committee (12/NTB/59), and parents provided written informed consent.

### Consent for publication

Not applicable.

### Availability of data and materials

The datasets generated and/or analysed for the performance of RBV are available in the RBV repository, https://github.com/whitneywhitford/RBV.

### Authors’ contributions

All authors were involved in the concept design and refinement of RBV. WW developed and tested the software, and wrote the manuscript. JCJ critically reviewed the manuscript. All authors read and approved the final manuscript.

## Acknowledgements

The author(s) wish to acknowledge the contribution of NeSI high-performance computing facilities to the results of this research. NZ’s national facilities are provided by the NZ eScience Infrastructure and funded jointly by NeSI’s collaborator institutions and through the Ministry of Business, Innovation & Employment’s Research Infrastructure programme. URL https://www.nesi.org.nz.

## Availability and Requirements

Project name: RBV

Project home page: https://github.com/whitneywhitford/RBV

Operating system(s): Linux

Programming language: Python 2.7

Other requirements: SAMtools 1.3 or higher, tabix

License: GPL v3

Any restrictions to use by non-academics: None

